# AFsample: Improving Multimer Prediction with AlphaFold using Aggressive Sampling

**DOI:** 10.1101/2022.12.20.521205

**Authors:** Björn Wallner

## Abstract

The AlphaFold neural network model has revolutionized structural molecular biology with unprecedented performance. We demonstrate that by stochastically perturbing the neural network by enabling dropout at inference combined with massive sampling, it is possible to improve the quality of the generated models. We generated around 6,000 models per target compared to 25 default for AF2-multimer, with v1 and v2 multimer network models, with and without templates, and increased the number of recycles within the network. The method was benchmarked in CASP15, and compared to AF2-multimer it improved the average DockQ from 0.41 to 0.55 using identical input and was ranked at the very top in the protein assembly category when compared to all other groups participating in CASP15. The simplicity of the method should facilitate the adaptation by the field, and the method should be useful for anyone interested in modelling multimeric structures, alternate conformations or flexible structures.

**Availability:** AFsample is available online at *http://wallnerlab.org/AFsample*.

## 1 Introduction

The unprecedented accuracy of AlphaFold (Jumper *et al*., 2021) has transformed the field of computational and structural biology. It is now possible to achieve highly accurate predictions on par with experimentally determined structures. AlphaFold has rapidly become the go-to method for protein structure prediction for both monomers and multimer prediction pipelines (Cramer, 2021; Terwilliger *et al*., 2022).

A key to the success of AlphaFold is its ability to assess the accuracy of its own predictions. AlphaFold estimates the per-residue accuracy using the predicted LDDT (Mariani *et al*., 2013) (pLDDT). In addition, it also calculates a predicted TMscore (Zhang and Skolnick, 2004) (pTM) from the predicted aligned error (PAE) matrix (Jumper *et al*., 2021). The correlation for pLDDT and pTM to its actual values are 0.76 and 0.85, respectively (Jumper *et al*., 2021), and this correlation is maintained even for high-quality predictions. For multimer prediction, an inter-chain predicted TMscore (ipTM) is calculated from the PAE of the inter-chain distances (Jumper *et al*., 2021).

Given enough evolutionary-related sequences, AlphaFold predicts monomers with very high accuracy even without using structural templates (Jumper *et al*., 2021). However, for multimers, this is not necessarily true (Bryant *et al*., 2022), the evolutionary signal constraining multimers is much weaker, and for these cases, more sampling might improve the prediction. The need for more sampling was also the reason why the default number of sampled models in AlphaFold-multimer was increased from 5 in version 1 (v1) to 25 in version 2 (v2) (Evans *et al*., 2021). Furthermore, predicting transient interactions or interactions with flexible binding partners, such as short peptides or disordered regions, requires even more sampling to achieve optimal performance (Johansson-Åkhe and Wallner, 2022).

For complex cases, simply increasing the number of sampled models might not be enough if the evolutionary constraints have trapped the prediction in a local minimum in the conformational landscape (Roney and Ovchinnikov, 2022) or the if the evolutionary constraints are weak. In such cases, increasing the number of times the prediction is recycled in the network can improve performance (Mirdita *et al*., 2022). Another option is to randomly perturb or alter the input MSA, which has been shown to enable better sampling of the conformational landscape and prediction of multiple conformational states (Alamo *et al*., 2022; Wayment-Steele *et al*., 2022). An alternative way to achieve more diversity among the generated models is to enable the dropout layers in the neural network (Johansson-Åkhe and Wallner, 2022; Mirdita *et al*., 2022). The dropout layers in a neural network are commonly used only when training the networks to cause them to learn different redundant solutions to the same problem by stochastically dropping some of their weights (putting them to zero). The dropout rate in the AlphaFold network is 10%-25%, depending on the network module. By activating these layers at inference, the network will naturally sample the uncertainties (Gal and Ghahramani, 2016) in the structure prediction, and the structural diversity of the sampled models will be increased.

## 2 Results

Here, we present AFsample that significantly improves over AlphaFold2-multimer (AF2-multimer) baseline. The method was the most successful in CASP15 for multimer prediction (Figure 1a). It improved the average DockQ (Basu and Wallner, 2016) from 0.41 for AF2-multimer (NBIS-AF2-multimer) to 0.55, with eight targets showing considerable improvements with more than 0.4 DockQ units, essentially going from incorrect to high-quality predictions (Figure 1b).

**Figure 1:**
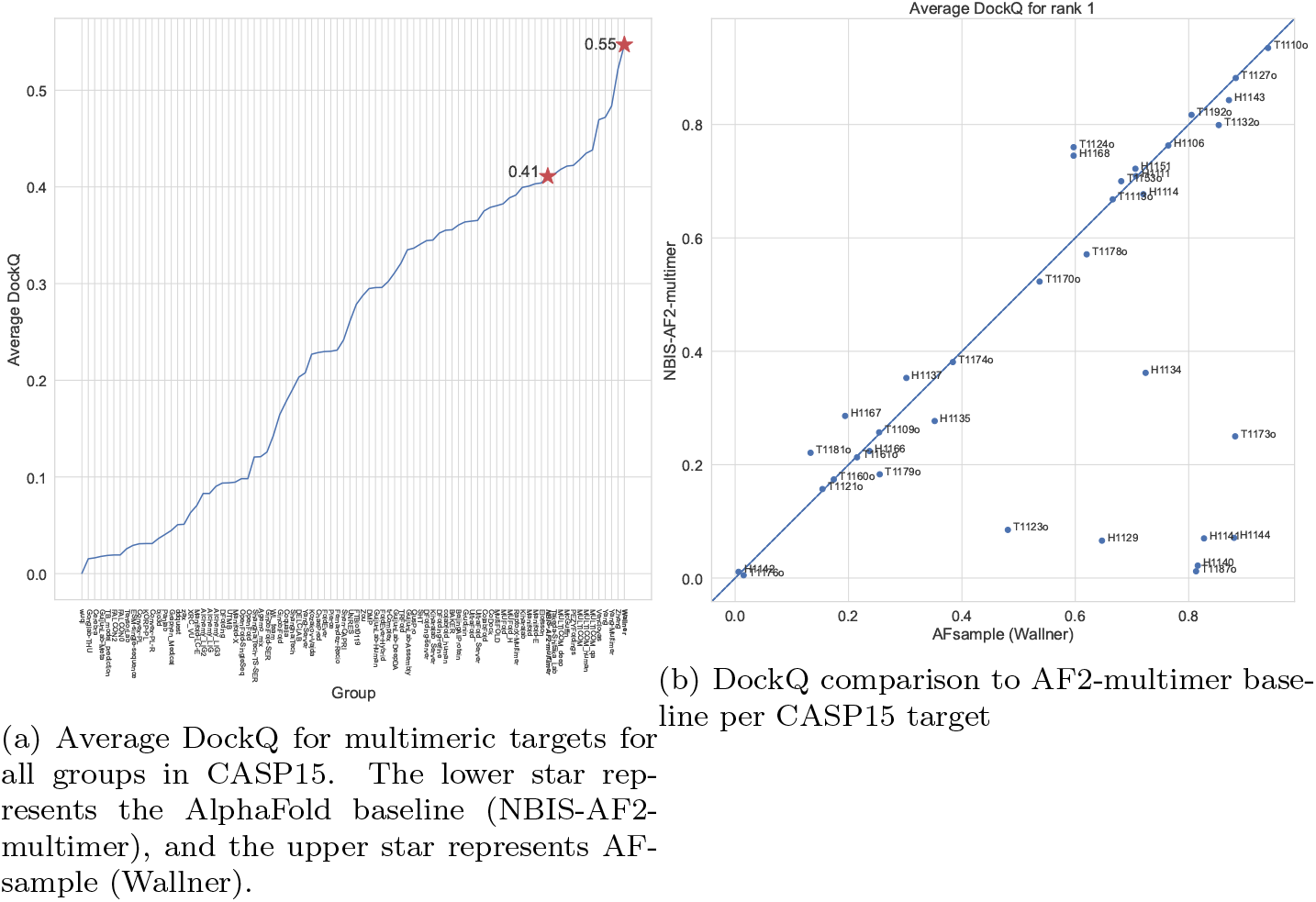
AFsample (Wallner) performance on common CASP15 multimer target compared AlphaFold baseline (NBIS-AF2-multimer).

In short, AFsample generates a large pool of models using different settings (Table 1). The self-assessment ranking score (`ranking_confidence`), which for multimer is a linear combination of the predicted interface TMscore (ipTM) and the predicted TMscore (pTM), 0.8ipTM + 0.2pTM, is used for selection. Six different settings were used, and they all involved using dropout to increase the model diversity. Both `v1` and `v2` of the multimer neural network weights and increased number of recycles were also utilized to increase the diversity further. The number of recycles for `v1` and `v2` were optimized in a previous study of peptide-protein predictions (Johansson-Åkhe and Wallner, 2022). Four settings involved selective dropout, where we only activated dropout in the Evoformer part of the network and not in the structural module. Despite the name, the bulk of structure prediction in AlphaFold is not performed in the structural module but in the much more extensive Evoformer network. In a previous study, we observed an improved correlation between `ranking_confidence` and actual DockQ using selective dropout with no dropout in the structural module (Johansson-Åkhe and Wallner, 2022). For each setting, on the order of 1,000 models were generated per setting for a total of 6,000 models per target.

**Table 1:** Different settings of AlphaFold used in AFsample, *Version* refers to the version of the multimer neural network weights, *Templates* refers to if structural templates were used or not, *Dropout* refers to if dropout was enabled, *Recycles* refers to how many recycles was used (default 3). ^*^No dropout in structural module

| Version | Templates | Dropout | Recycles |
| --- | --- | --- | --- |
| multimer_v1 | Yes | Yes | 3 |
| multimer_v1 | No | Yes* | 3 |
| multimer_v1 | No | Yes* | 21 |
| multimer_v2 | Yes | Yes | 3 |
| multimer_v2 | No | Yes* | 3 |
| multimer_v2 | No | Yes* | 9 |

It is clear the sole reason for the improved performance of AFsample is improved sampling as AFsample was using identical MSAs as the AF2-multimer baseline. The need for sampling is illustrated further by target H1144 (Figure 2). Target H1144 is a nanobody interaction, nanobodies and regular antibody interactions are particularly challenging to model since the interaction with the antigen is highly specific, and the exact shape and location of the epitopes can vary significantly depending on the antigen (Chiu *et al*., 2019). Here, only three out of 6,000 models obtained a `ranking_confidence`>0.8, out of which all were of high quality with a `DockQ`>0.8. In fact, only five high-quality models could be found in the whole set of 6,000 models sampled. Thus, without substantial sampling, most likely, no high-quality model would have been generated at all.

**Figure 2:**
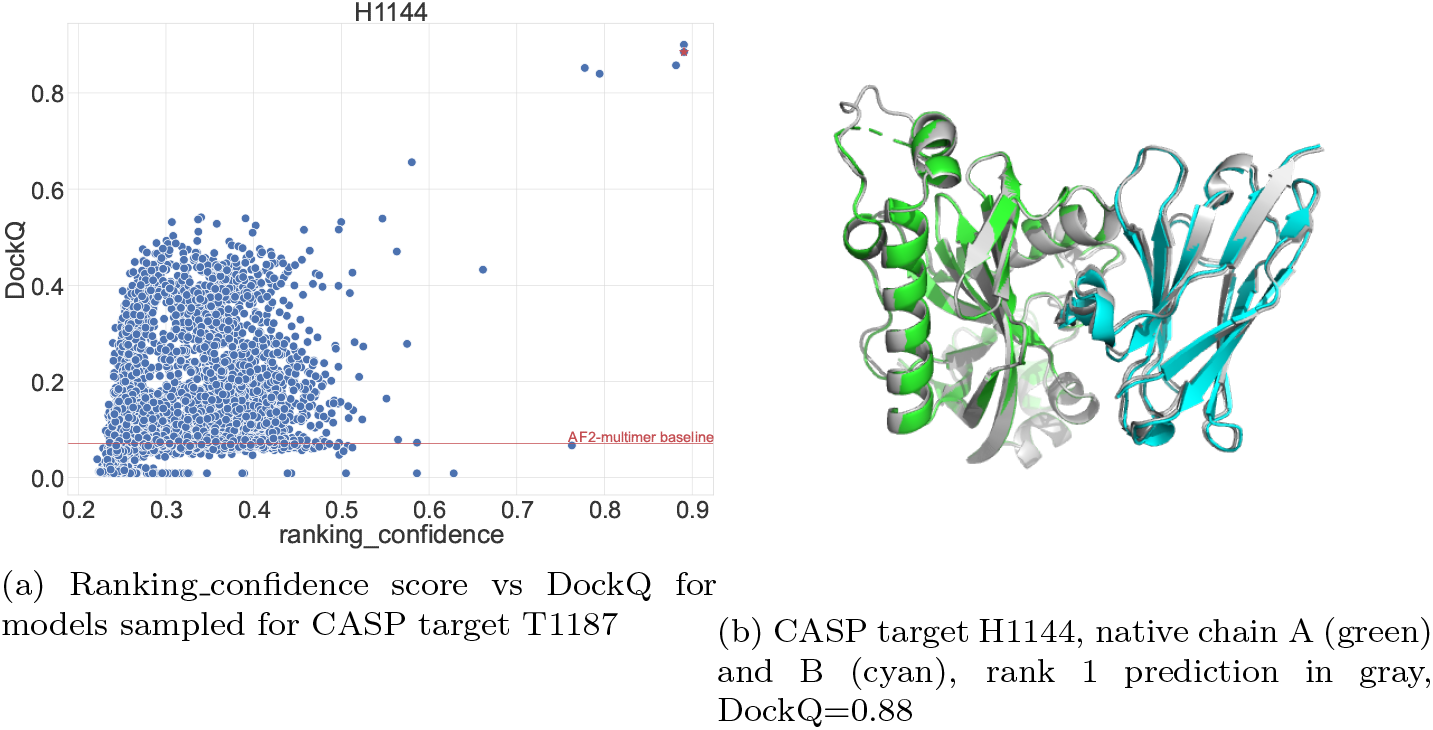
Example to illustrate the sampling process from CASP15

The other example is T1187o (Figure 3). Target T1187o is a dimer of the Uniprot ID `Q94EW1`. The AlphaFold prediction of the monomer is almost perfect (https://alphafold.ebi.ac.uk/entry/Q94EW1), but the model of the dimer is completely wrong with a DockQ very close to zero (Figure 2a). However, with improved sampling, it is possible to generate several high-quality models (38/6000). Still, selecting the best possible model was not straightforward as one-third of the models with a `ranking_confidence`>0.8 actually had a wrong domain orientation indicated by the low DockQ score.

**Figure 3:**
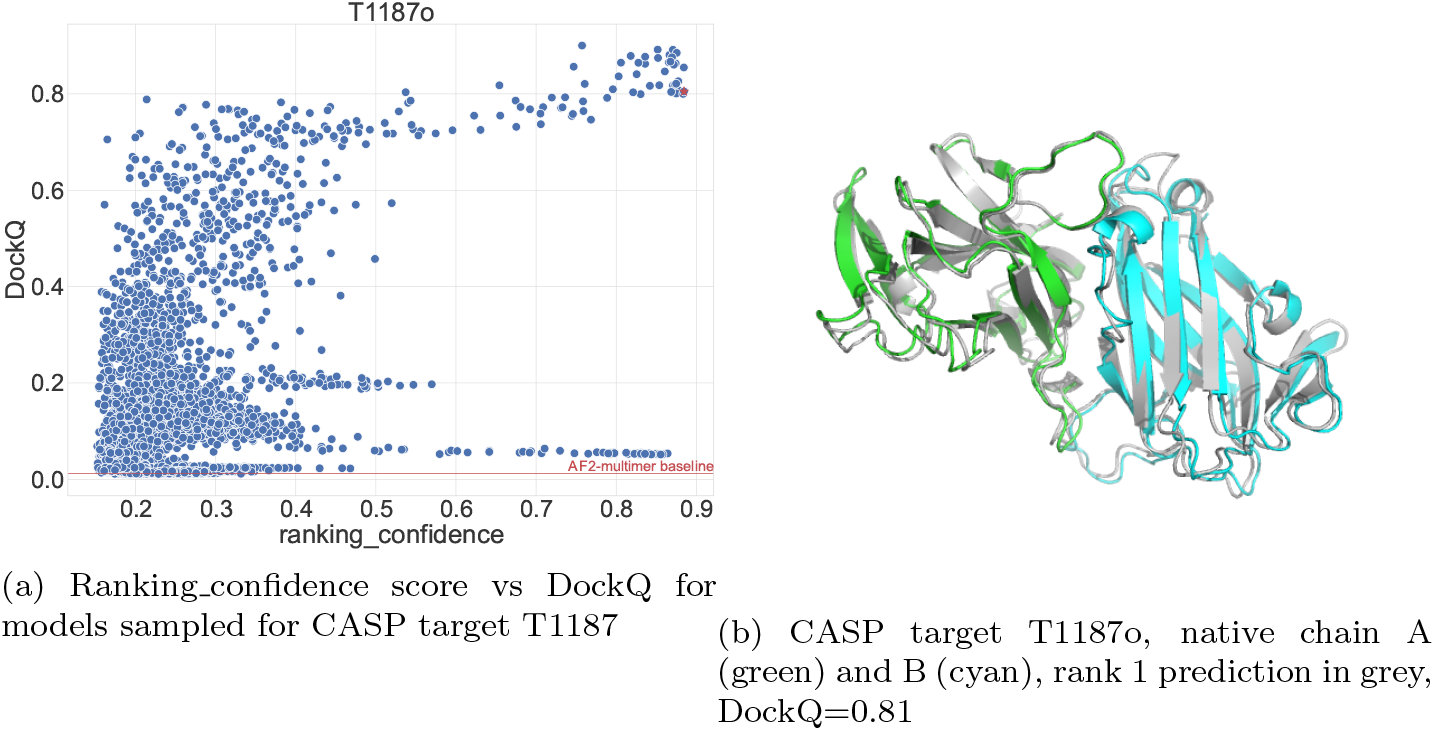
Examples from CASP15

AFsample requires more computational time than AF2-multimer baseline, as it generates 240x models, and including the extra recycles, the overall timing is around 1,000x more costly than the baseline. Thus, a good strategy is first to run the AF2-multimer baseline prediction and only use more sampling if needed. A good criterion for this is to use a `ranking_confidence`>0.8 and require at least a couple of structures at this level of confidence. Finally, given that AlphaFold inference in many cases is relatively fast, a minute or less per model, a couple of days on a single GPU to get high-quality results for a difficult target is not an enormous amount of computational time.

The results from CASP15 demonstrate that the best way to model multimeric protein assemblies today is to use AFsample. However, it is very likely that sampling will improve the performance of AlphaFold in any setting. Dynamics, flexibility, or simply the sheer complexity of large molecular assemblies will all require more sampling. The success of sampling relies heavily on the excellent internal scoring function in AlphaFold, which so far has proven to be exceptionally good.

## 3 Methods

AFsample is available as a command line interface using a modified version of the official AlphaFold release. The modified version is streamlined to produce many models, contains functionality to parallelize the computations to independent jobs, and exposes several internal parameters of AlphaFold to allow use it at its full potential. A script `run_afsample.sh` is provided that reproduces the method that participated in CASP15 under the name `Wallner`.

### Filter the output

To enable the generation of thousands of models, the amount of data saved per model was limited to the predicted aligned error matrix (PAE), per residues predicted LDDT (`pLDDT`), overall predicted TMscore(`pTM`), predicted interchain TMscore (`ipTM`), and `ranking_confidence`. This reduces the size of the data saved per model by approximately a factor 100.

The flag `--output_all_results` will restore the default behavior and output all data structures.

### Checkpointing

Checkpointing is made default, it will not recalculate MSAs if they exist and if a model exists, it will continue to the next model.

### Added functionality

The most important modifications and exposed functionalities are described below:

`--model_preset` Modified to allow using both v1 and v2 models, the following presets are allowed: `multimer_v1, multimer_v2, multimer` (defaults to v2), `multimer_all` (both v1 and v2)
`--nstruct` the number of structures to output, works both for monomer and multimer protocols and replaces the `--num_multimer_predictions_per_model`, which was exclusive to the multimer protocol.
`--max_recycles` the number of times a prediction is recycled in the neural network, default is 3.
`--dropout` enables the dropout layers at inference.
`--dropout_structure_module` if `False`, it will not have the dropout layers enabled in the structure module.
`--suffix` option to add a descriptive suffix to each model name to enable output to the same output folder.
`--no_templates` do not use any templates, faster than filter by date, which requires parsing all hits.
`--seq_only` will only run the sequence searches to create the MSAs and template hits, which is useful for preparing input files for larger runs.
`--input_msa` option to input a multiple sequence alignment in STOCKHOLM format.
`--nstruct_start` which structure to start with, useful to split a large job into many smaller by using `--nstruct_start 20` and `--nstruct 21` it will create models `20` and `21`.
`--models_to_use` option to specify which neural network models from `model_preset` to use. i.e `model_1_multimer_v2, model_3_multimer_v2` will only run model 1 and model 3 from `multimer_v2`.

### Databases

Sequence databases were downloaded on April 22, 2022, and the PDB was updated May 2, 2022, using the download scripts (https://github.com/deepmind/alphafold/scripts/ provided by DeepMind. The following versions were used:

- Uniclust30 (Mirdita *et al*., 2017) version: `UniRef30 2021 03`
- Uniref90 (Suzek *et al*., 2015) from April 22, 2022.
- Uniprot, TrEMBL+SwissProt, from April 22, 2022.
- BFD database (Steinegger and Söding, 2018) `bfd metaclust clu complete id30 c90 final seq.sorted opt cs219.ffindex` `MD5 hash: 26d48869efdb50d036e2fb9056a0ae9d`
- Mgnify version: 2018 12
- PDB from May 2, 2022.

### All-atom relaxation

In regular AlphaFold, each model is constrained relaxed in the Amber99sb force field (Hornak *et al*., 2006) using openMM (Eastman *et al*., 2017). To save computational time, the all-atom relaxation step is skipped for each model. Instead, the `run_relax_from_results_pkl.py` script is provided, which performs the relaxation step for a given result `pickle` as outputted by AlphaFold. Since none of the scores depends on this step, the relaxation can be performed only for a smaller subset of the models that are high scoring or are selected by some other criteria.

### Benchmark

In the CASP15 benchmark, AFsample (`Wallner`) and AF2-multimer baseline were run with exactly the same multiple sequence alignments (MSAs) and templates. The alignments were created with the large database setting: `--db_preset=full_dbs` using the AF2-multimer baseline server (`NBIS-AF2-multimer`). They were made available by the CASP organisers, and these were the MSAs used by the `Wallner` group in CASP15. The DockQ (Basu and Wallner, 2016) scores for all methods that participated in CASP15 were downloaded from the CASP15 website. In the case of multiple interfaces, DockQ is calculated for each interface and then averaged. The rank 1 models from each method were used to calculate the average DockQ for the multimer targets for each method.

## Data Availability

The MSAs and template information used in the CASP15 benchmark are available here: http://bioinfo.ifm.liu.se/casp15/

## Code Availability

AFsample is free, open-source software (Apache) and is available from here: http://wallnerlab.org/AFsample

## Acknowledgments

This work was supported by a WASP-DDLS grant from Knut and Alice Wallenberg (KAW), Swedish Research Council grant, 2020-03352, The Swedish e-Science Research Center, and Carl Tryggers stiftelse för Vetenskaplig Forskning, 20:453. The computations were performed on resources provided by the Swedish National Infrastructure for Computing (SNIC), Knut and Alice Wallenberg (KAW) (Berzelius), and Linköping University (LiU) at the National Supercomputer Center (NSC) in Linköping.

